# Temporal dynamics of cognitive flexibility in adolescents with anorexia nervosa: a high density EEG study

**DOI:** 10.1101/2022.03.24.485666

**Authors:** Cristina Berchio, Lucie Clémentine Annen, Ynès Bouamoud, Nadia Micali

**Affiliations:** Department of Psychiatry, Faculty of Medicine, University of Geneva, Geneva, Switzerland; Department of Pediatrics, Gynecology and Obstetrics, Faculty of Medicine, University of Geneva, Geneva, Switzerland; Great Ormond Street Institute of Child Health, University College London, London, UK

**Keywords:** Anorexia nervosa, adolescence, cognitive flexibility, ERP, source imaging

## Abstract

Cognitive rigidity is a core symptom of anorexia nervosa (AN) and is associated with treatment resistance. Nevertheless, studies on the neural basis of cognitive flexibility (CF) in adolescent AN are rare.

The aim of this study was to investigate temporal brain dynamics of CF in adolescents with AN and their tendency to experience anxiety. To address this aim, participants performed a Dimensional Change Card Sorting task during high-density EEG recording. Transient state and stable traits of anxiety were measured with the State-Trait Anxiety Inventory. Data were collected on 22 girls with AN, and 23 healthy controls (HC) (aged 12-20 years old). Evoked potentials were investigated using microstates analysis. Effects of trait and state anxiety on microstates parameters were investigated performing correlations across groups and in the AN group only.

Adolescents with AN did not differ from HC with respect to behavioral switching cost, despite a greater overall accuracy. Adolescents with AN showed altered visual orienting processes (N100 micro-state), and earlier attentional processing to task switching (P300b microstate) when compared to HC. Reduced activation in a network involving the pre-central cortex, the cerebellum, and the posterior cingulate was identified in adolescents with AN. Trait and state anxiety were correlated with atypical AN microstates across groups.

The present findings suggest CF manifests with inefficient behavioral and neural strategies in adolescents with AN, at early stages in the illness. Additionally, this study provides evidence that transient and stable tendencies to experience anxiety could represent a vulnerability to developing CF impairments in adolescent AN.

## 1. Introduction

Anorexia nervosa (AN) is a mental illness characterized by body image disturbance and behaviors that interfere with weight gain (A.P.A., 2013). Behavioral rigidity and psychological inflexibility are associated with AN(Buzzichelli, Marzola, Amianto, Fassino, & Abbate-Daga, 2018), and have an impact on symptoms and treatment resistance(Tchanturia, Lloyd, & Lang, 2013; Treasure & Schmidt, 2013).

Behavioral and mental rigidity in AN have been investigated by examining cognitive flexibility (CF)(Miles, Gnatt, Phillipou, & Nedeljkovic, 2020). Cognitive flexibility (CF) is the ability to adapt one’s mental strategies to unexpected situations and environmental changes, and it crucially depends on attentional control (Canas, Fajardo, & Salmeron, 2006). Several studies have documented CF impairments in adults diagnosed with AN (Miles et al., 2020; Roberts, Tchanturia, Stahl, Southgate, & Treasure, 2007; Roberts, Tchanturia, & Treasure, 2010; Tchanturia et al., 2012; Westwood, Stahl, Mandy, & Tchanturia, 2016).

Deficits in adolescents with AN require further investigation. Few studies have documented worse performance in flexibility tasks between adolescents with AN and healthy controls (HC) (Lang et al., 2015; McAnarney et al., 2011), while others failed to find significant group differences (Andrés-Perpiña et al., 2011; Bohon, Weinbach, & Lock, 2020; Dmitrzak-Weglarz et al., 2011; Fitzallen, Sagar, Taylor, & Bora, 2021; Van Autreve et al., 2016).

Neuroimaging studies of CF in AN are scarce, especially in adolescence. Findings in adult populations provide evidence that dysfunctional frontal-parietal control networks might underlie CF difficulties in AN (Lao-Kaim et al., 2015; Sato et al., 2013; Zastrow et al., 2009). A recent study using a visuo-spatial shifting task, with shapes and colors as stimuli, documented reduced activations in adolescents with AN, compared to controls, in the inferior and middle occipital lobe, the lingual regions, the fusiform cortex, and the cerebellum (Castro-Fornieles et al., 2019). The few available studies also seem to suggest that, during adolescence, brain alterations may be reversible after treatment/weight gain (Bohon et al., 2020; Castro-Fornieles et al., 2019). However, it remains unclear whether behavioral and neural deficits of CF could be explained by other clinical traits interacting with the active state of the illness.

Individuals with high trait anxiety tend to have worse performance in tasks involving CF(Wilson, Nusbaum, Whitney, & Hinson, 2018). Previous work has indicated a negative correlation between behavioral CF performance (mistakes, reaction times) and anxiety measures (Han et al., 2011). Neuroimaging evidence indicates that trait anxiety induces reduced control of the prefrontal cortex during conflict tasks (Bishop, 2009). Electrophysiological data have also shown that induced anxiety is associated with enhanced attention allocated to stimuli, and with impaired cognitive-behavioral flexibility(Cornwell, Mueller, Kaplan, Grillon, & Ernst, 2012).

There is good evidence that both state and trait anxiety are associated with AN (Raney et al., 2008; Schulze, Calame, Keller, & Mehler-Wex, 2009). Although the behavioral literature, especially in adults with AN, seems to exclude the impact of depression and anxiety on CF dysfunction (Fuglset, 2021), less is known about anxiety in adolescents with AN. Therefore, whether trait- or state anxiety contributes to CF (at the behavioral and neural level) in adolescents with AN also requires further investigation.

High-density electroencephalography (EEG) and event-related potentials (ERP) allow an investigation of behavioral responses and large-scale networks with millisecond precision(Michel & Murray, 2012; Murray, Brunet, & Michel, 2008). High density ERP are therefore capable of capturing brain dynamics of CF. Furthermore, this method is non-invasive, making it particularly suitable for the study of childhood and adolescent psychiatric disorders(Berchio & Micali, 2022).

Early visual ERP components are believed to reflect sensory and perceptual processing (i.e., P100, N100 (Woodman, 2010)). P100, the first visual component, is usually localized in the middle occipital cortex, while the subsequent negative wave in the parietal lobe (Creel, 2019).

An ERP component that is usually linked to CF is the P300 (Barcelo & Cooper, 2018; Gajewski & Falkenstein, 2011; Kopp, Steinke, & Visalli, 2020). The P300 occurs roughly 300–700 milliseconds (ms) post-stimulus onset, with a broad central-parietal scalp distribution (Woodman, 2010). The P300 a and the P300 b are sub-components of the P300, which are believed to reflect attentional and memory processes, respectively(Ghani, Signal, Niazi, & Taylor, 2020). Brain sources of the P300 a are estimated mainly in the prefrontal cortex, of the P300 b in parietal and inferior temporal regions (Bledowski et al., 2004).

To date, high density EEG has not been used to study CF in adolescents with AN. High density ERPs may provide novel insights into brain correlates of CF in adolescent AN.

The main objective of the present study was to evaluate neural activity assessed by high density EEG during a CF task in adolescents with AN compared to a group of HC. Furthermore, we aimed to assess whether trait or state anxiety influenced CF both at the neural and behavioral level.

We hypothesized that we would observe impaired brain temporal dynamics in AN adolescents compared to HC during task switching. Since the P300 is a marker of CF, we expected that adolescents with AN would have P300 deficits, mainly in frontal-visual networks. We expected higher scores of trait-state anxiety in AN when compared to HC, and we hypothesized that brain network characteristics of CF would be predicted by trait and state anxiety. We expected worse CF in AN (i.e., slower reaction times, more perseverative errors), compared to HC, and that higher levels of trait and state anxiety would be associated with behavioral impairments.

## 2. Methods

### 2.1. Participants

Twenty-two girls with AN and twenty-three HC participants were included in the study. All participants were females, and their age ranged from 12 to 20 years (We choose to refer to our population as adolescents according to mean ages of the samples and based on indications from literature (Sawyer, Azzopardi, Wickremarathne, & Patton, 2018)).

Patients with AN were recruited from the adolescent and adult ED services at Geneva University Hospital (HUG). Individuals received diagnoses of restricting type AN (n=19), and binge/purge type AN (n=4) by the respective multidisciplinary teams during clinical evaluations.

Controls were recruited through advertisements posted in local community centers, universities, and schools. General exclusion criteria were history of any neurological disorder or brain damage; absence of any psychopathology and ED symptoms.

All controls were screened for psychopathology using the Strengths &Difficulties Questionnaire (SDQ) [completed by parents and youths for participants aged 12-17 years old, and youths only for participants > 17 years old](Goodman, 1997). Furthermore, ED symptomatology was assessed in 17 controls using the Eating Disorder Examination Questionnaire (EDE-q) (Fairburn & Beglin, 1994)[a 28-item self-report questionnaire assessing eating disorder features; 4 subscales (Restraint, Eating Concern, Shape Concern and Weight Concern) and a global score are obtained], and in 6 by the caregiver filling in the ED section of The Development and Well-Being Assessment (DAWBA) questionnaire (Goodman, Ford, Richards, Gatward, & Meltzer, 2000). All patients with AN completed the EDE-q.

Participants also completed the State–Trait Anxiety Inventory according to their age (STAI; adult version [(Spielberger, 1983) (ages 16 through 20)] or child versions [(Turgeon & Chartrand, 2003) (ages 12 through 16)]. For subsequent analysis, scores were z-transformed using normative scores of adults and children). Information about participants’ ethnicity, family education was also collected.

During the first visit, cognitive functioning was assessed for all participants using 4 sub-scales of the Wechsler Intelligence Scale for Children Fifth Edition[WISC-V, ages 12 through 16, (Wechsler, 2014)] or the Wechsler Adult Intelligence Scale | Fourth Edition [WAIS-IV, ages 16 through 20, (Wechsler, 2008)]: similarities, vocabulary, matrix reasoning, and block design. These subscales were administered to ensure no confounding due to cognitive function.

None of the controls meet problems in any domain. Two adolescents with AN were diagnosed as having an anxiety disorder, one was diagnosed with depression: one of these was taking psychotropic medication (Selective Serotonin Reuptake Inhbitor (SSRI)).

Participants who were aged 16 and older, and the parents of those below the age of 18, provided written informed consent. The research was conducted according to the principles of the Declaration of Helsinki and approved by the University of Geneva research ethics committee. Participants received vouchers from a general store as a thank you for their participation in the study.

### 2.2 Experimental paradigm

In the DCCS (Zelazo, 2006), participants are required to sort bivalent test cards using rules. In this study, we applied a version of the DCCS that has been proven to be suitable for older children and neuroimaging data collection [see,(Morton, Bosma, & Ansari, 2009)]. Two bivalent images were always presented at the bottom of the screen: a red rabbit and a blue truck. On each trial, a fixation cross (750 ms) was followed by an instruction period (1750 ms), and a response period (2000 ms). After the fixation period, participants were instructed on the trial rule (“s” for shape or “c” for color), which was followed by the target stimulus: a blue rabbit or a red truck. The target stimulus matched each image presented at the bottom on a single dimension (either color or shape). Participants responded by pressing one of two arrows on a keyboard using the index and middle-finger of their right hand. By pressing the right arrow, participants sorted the stimulus to the location of the right target (i.e., the blue truck); by pressing the left arrow they sorted the stimulus to the location of the left target (i.e., the red rabbit). Trials were separated by a 750 ms inter-trial interval.

Trials were administered in 2 blocks of 160 trials. Switch trials (n=80) consisted of switch sorting rule trials (e.g., “s”, “r”), and repeat trials (n=80) of sorting rule repeated (e.g., “c”, “c”). Trials were administered in a semi random order in each block.

To ensure participants understood the task, the experiment was preceded by a training block (10 trials). Total task duration was approximately 20 minutes.

### 2.3 EEG data acquisition and pre-processing

EEG data were acquired at 1000 Hz using a 256-channel system (EGI, Philips Electrical Geodesics, Inc.). Electrode impedance was kept below 30 kΩ, and data were acquired with the vertex (Cz) as reference electrode.

EEG data time-locked to target stimuli onset (respond period) were averaged separately for repeated and switching trials. EEG data were averaged only for correct trials. Data were band-pass filtered between 0.4 and 40 Hz, and a 50-Hz notch filter was applied. The original montage was reduced from 256 to 204 channels to exclude bad channels located at the periphery (see(Berchio, Rodrigues, Strasser, Michel, & Sandi, 2019)). Infomax-based Independent Component Analysis (ICA) was applied to remove eye blink, eye-movements and electrocardiogram artifacts based on map topographies, and the time course of the ICA component. Epochs for ERP analysis began 100 ms before stimulus, and ended 600 ms after its onset. Epochs contaminated by artifacts (muscle, eye-blink/movements) were excluded by visual inspection. Epochs were re-referenced to the average, and down-sampled to 250 Hz. Data pre-processing was performed using the CARTOOL Software(Brunet, Murray, & Michel, 2011). There were no group differences in the number of trials accepted (AN: repeated trials: *M*= 74.36 *SD*=5.1, switching trials: *M*=73.90, *SD*=4.19; HC: repeated trials: *M*=71.34, *SD*=7.82, switching trials: *M*=72.65, *SD*=6.04; *p >* 0.14. two tailed-t test)).

### 2.4 Behavioral analyses

For each behavioral measure (Accuracy, Reaction times (RT), Errors), a repeated-measures ANOVA was conducted for comparisons between groups, with Trials as within-subject factor (‘Repeat’ vs. ‘Switch’) and Group as between-subject factor (‘AN’ vs. ‘HC’). To clarify main effects and interactions, post hoc analysis used paired t-tests with *p* < .05 as the significance threshold after a Bonferroni correction for multiple comparisons.

### 2.5. ERP analysis

#### 2.5.1 ERP microstate analysis

ERPs were analyzed using microstate analyses(Michel & Murray, 2012; Murray et al., 2008). This k-means cluster-based approach is a reference-free method that can be applied to classify ERP components, as well as experimental conditions and groups. The advantage of applying this method is that changes in map topographies are indicative of changes in brain network dynamics (Lehmann, 1987; Vaughan, 1982).

K-means clustering was first applied on the grand averages (two conditions, two groups) [for more details see also (Murray, De Lucia, Brunet, & Michel, 2009; Pascual-Marqui, Michel, & Lehmann, 1995)]. To make physiologically plausible assumptions, the maximum number of possible clusters was set to 20, and a temporal rejection criteria (minimum duration accepted equal to 16 ms) was also applied. Subsequently, the optimal number of clusters was identified based on the meta-criterion implemented in Cartool.

To statistically confirm the optimal model identified, the corresponding maps were reassigned to individual ERP data using the toolbox “Randomization Graphical User Interface” (Ragu)(Koenig, Kottlow, Stein, & Melie-García, 2011). This procedure consists of the assignment, at each moment in time, to one of these maps based on an estimated spatial correlation index [for technical details, see (Koenig, Stein, Grieder, & Kottlow, 2014; Rohde et al., 2020)].

The assignment procedure allows to extract microstates variables of interest. Variables investigated in this study were: frequency of occurrence (as in index of presence/absence), first offset of a given map (as a temporal parameter index), and the area under the curve ((AUC), as an additional global measure of occurrence).

To assess differences between groups, microstates variables were compared using appropriate statistical tests according to data distribution.

#### 2.5.2 Brain imaging analyses between groups

Network differences between groups were tested using a linear distributed inverse solution model, which considers each voxel as a possible source of activity (LORETA,(Pascual-Marqui, Esslen, Kochi, & Lehmann, 2002)). The inverse solution was estimated on average ERPs, using a Locally Spherical Model with Anatomical Constraints [LSMAC model (Brunet, Murray, Michel, & neuroscience, 2011)], and on 5018 voxels located on the grey matter of a brain template of the Montreal Neurological Institute (http://www.bic.mni.mcgill.ca/brainweb). Following the computation of the inverse solution, data were transformed by a modified z-score normalization implemented in Cartool (3.80) (for technical details see:(Michel & Brunet, 2019)).

To minimize the problem of multiple comparisons, brain network analyses were performed using a randomization test implemented in Cartool on 116 regions of interest (ROI, Automated Anatomical Atlas (Tzourio-Mazoyer et al., 2002), 116 ROIs), with a *p* value less than .05, and an average time window of interest. Student unpaired two-tailed *t* tests were used for post hoc analyses. Time-windows for brain-source analyses were determined based on statistical evidence of ERP microstate analysis.

#### 2.5.3 Associations between behavior, clinical scores, and microstates presence

In order to investigate relationships between performance on the DCCS task and anxiety, Spearman correlations were conducted between behavioral performance of task switching (Accuracy, RT, mistakes) and state-trait anxiety measures.

To explore potential anxiety correlates of between-group differences, correlations were performed across groups and in the AN group only. A point-biserial correlation was used to measure the strength and direction of the association between state/trait anxiety and microstates presence.

Additionally, in the AN group, point-biserial correlations were performed to assess the relationship between microstates occurrence, eating disorders symptoms, and body mass index (BMI). For all correlations, bootstrapped confidence intervals (95%) were calculated to assess the significance of the effects estimated.

## 3. Results

### 3.1 Demographic and clinical variables

### 3.2 Behavior

Accuracy, RT, and errors on switch and repeat trials (see Table 2) were compared using repeated measures Analysis of Variance (ANOVA), with Trial Type (switch, repeat) and Group (AN, HC) as within- and between-subjects factors respectively. We found an effect of Trial Type on accuracy [F(1, 43)=30.9, p<.001], RT [F (1, 21) = 34.2, p<.001], and errors [F (1, 21) = 34.2, p < .001], with responses on switching trials significantly less accurate, slower, and more error prone than responses on repeated trials (post hoc tests, all p_s_ < .001). Importantly, there was a main effect of Group on accuracy, indicating greater accuracy independently of Trial Type in adolescents with AN when compared to HC [F (1, 43) =4.43, p=.041].

**Table 1.**

|  | HC |  | AN |  | *p* |
| --- | --- | --- | --- | --- | --- |
|  | *Mean* | *SD* | *Mean* | *SD* |  |
| Number of participants | 23 |  | 22 |  |  |
| Mean age (SD) | 15,56 | 3,34 | 15,81 | 2,41 | n.s |
| Body mass index | 20,74 | 2,81 | 16,86 | 1,53 | <.001 |
| Ethnicity (caucasian) | 81.1% |  | 91.3% |  |  |
| Parental education (one caregiver) |  |  |  |  |  |
| Secondary education | 35% |  | 30% |  | n.s. |
| University studies | 67% |  | 70% |  | n.s. |
| STAI*: |  |  |  |  |  |
| State | -0,691 | 0,53 | 0,571 | 0,98 | <.001 |
| Trait | -0,477 | 0,56 | 0,488 | 1,04 | <.001 |
| Cognitive function (WISC/WAIS) |  |  |  |  |  |
| Block design | 14,65 | 11,53 | 11,41 | 2,75 | n.s |
| Similarities | 15,78 | 8,06 | 14,68 | 2,62 | n.s |
| Matrix reasoning | 12,17 | 5,04 | 11,31 | 2,58 | n.s |
| Vocabulary | 15,65 | 9,27 | 13,13 | 2,66 | n.s |
| *z-scores |  |  |  |  |  |

**Table 2.** Behavioral datas.

|  | HC |  | AN |  |
| --- | --- | --- | --- | --- |
|  | *Mean* | *SD* | *Mean* | *SD* |
| **Accuracy (%):** |  |  |  |  |
| Switch trials | <u>92,42</u> | 6,37 | <u>94,36*</u> | 4,01 |
| Repeat trials | <u>96,42</u> | 2,89 | <u>97,13*</u> | 1,75 |
| **Reaction times (ms):** |  |  |  |  |
| Switch trials | 787,87 | 183,45 | 720,57 | 170,51 |
| Repeat trials | 730,09 | 183,46 | 685,61 | 133,13 |
| **Mistakes (%):** |  |  |  |  |
| Switch trials | 7,86 | 6,98 | 6,12 | 5,5 |
| Repeat trials | 2,8 | 2,55 | 2,65 | 3,21 |

\*Significant Main Group effect $p<.05$

### 3.3. ERP

Visual inspection of grand averages revealed the following ERP components to target stimuli (see Figure 2, A): the P100, N100, P200, P300a, P300b, and a late positivity.

**Figure 1.**
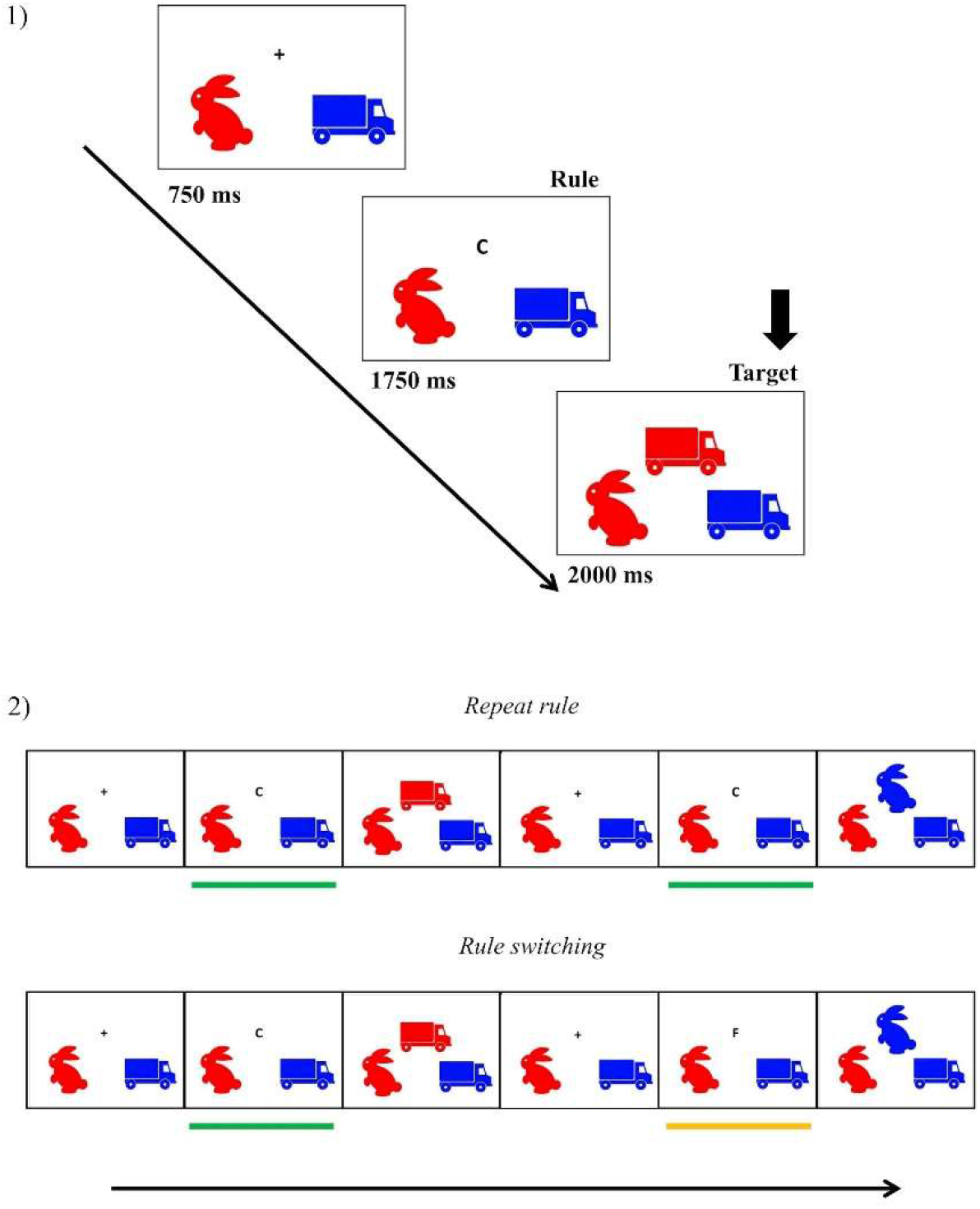
Schematic representation of the DCCS task. Each ‘rule’ (color or shape) is presented for 1750 ms, followed by a stimulus target (‘red truck’ or ‘blue rabbit’) for 2000 ms. Participants were asked to answer according to the rule of the trial.

**Figure 2.**
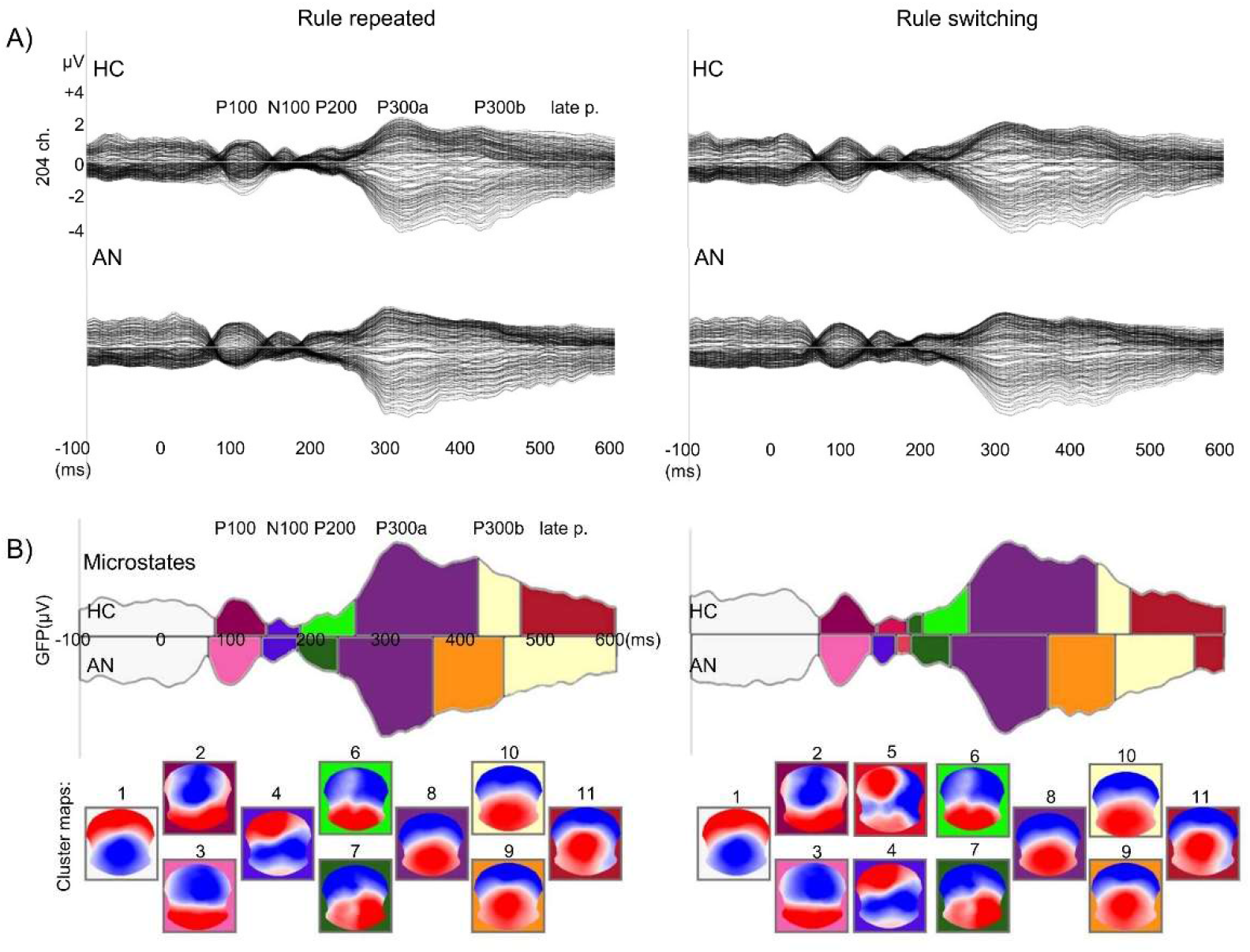
A) Grand averages (butterfly montage) of rule repeated and rule switching trials: adolescents with anorexia nervosa (AN), and healthy controls (HC). B) Results of k-means clustering on grand averages. Different colors highlight different classes of microstates. For each microstate, the corresponding EEG map is shown.

K-means clustering on grand averages identified eleven different map clusters: one for the baseline (map 1), two for the P100 (map 2, map 3), two for the N100 (map 4, map 5), two for the P200 (map 6, map 7), one for the P300a (map 8), two for the P300 b (map 9, map 10), and one for a late positivity (map 11).

For repeated trials, k-means clustering indicated potential differences between groups for the P100 (map 2 vs. map 3), P200 (map 6 vs. map 7), P300 a (map 8 latencies), P300 b (map 9 vs. map 10), and at a late positivity (map 11 latencies).

For switching trials, it indicated potential differences between groups at the P100 (map 2 vs. map 3), N100 (map 5 vs. map 4), P200 (map 6 vs. map 7), P200 (map 6 vs. map 7), P300 a (map 8 latencies), P300 b (map 9 vs. map 10), and at a late positivity (map 11).

To test whether group differences were statistically meaningful, the 11 maps were fitted back to the individual ERP data. The assignment procedure allowed to extract microstates parameters of interest: frequency of occurrence, offset, and the AUC. For all experimental conditions, individual data are plotted in Figure 3.

**Figure 3.**
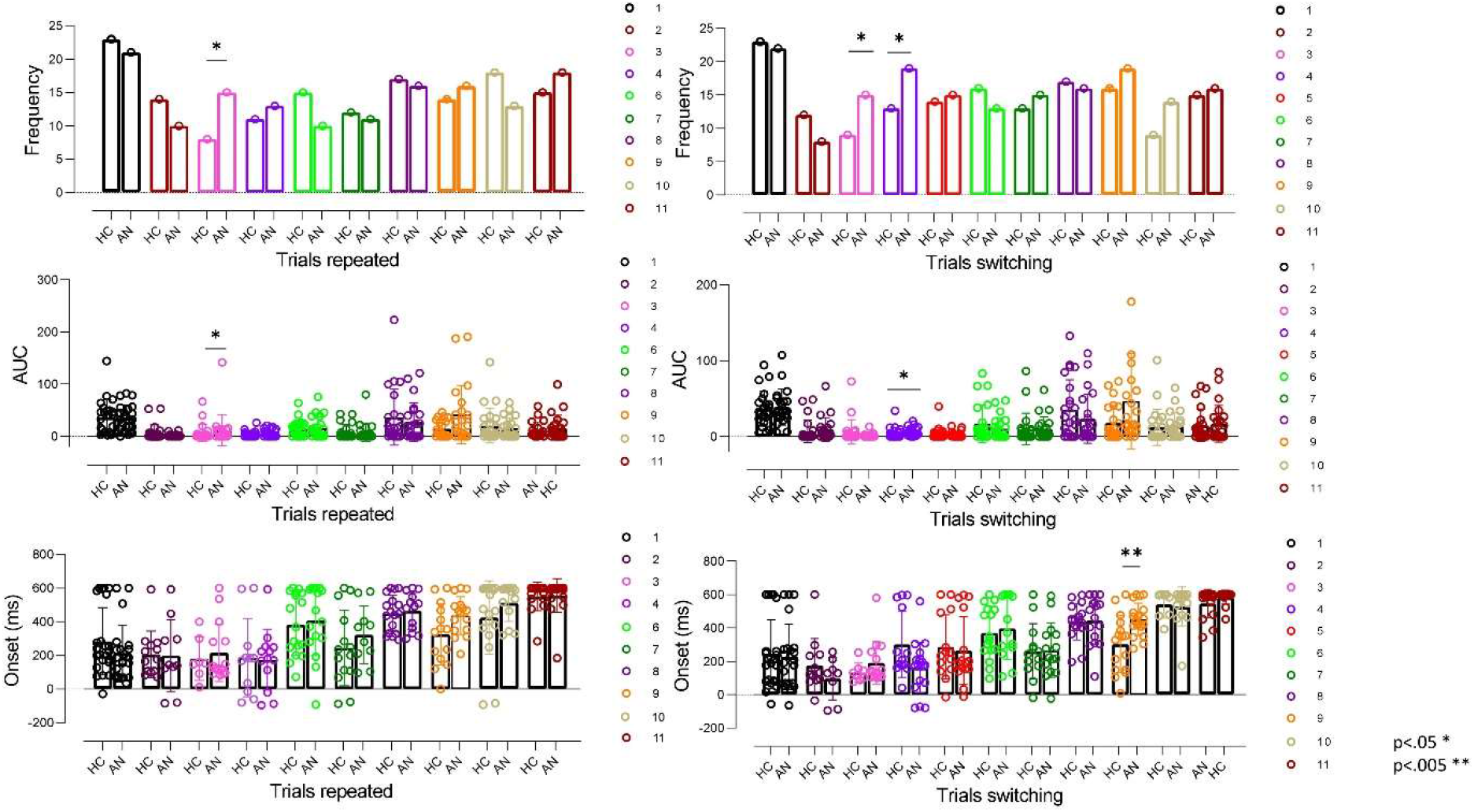
Microstates parameters of interest: frequency of occurrence, offset, and area under the curve (AUC). For all experimental conditions, individual data is plotted. Statistically significant differences between groups are marked by asterisks (\**p* <.05; \*\**p*<.005).

**Figure 4.**
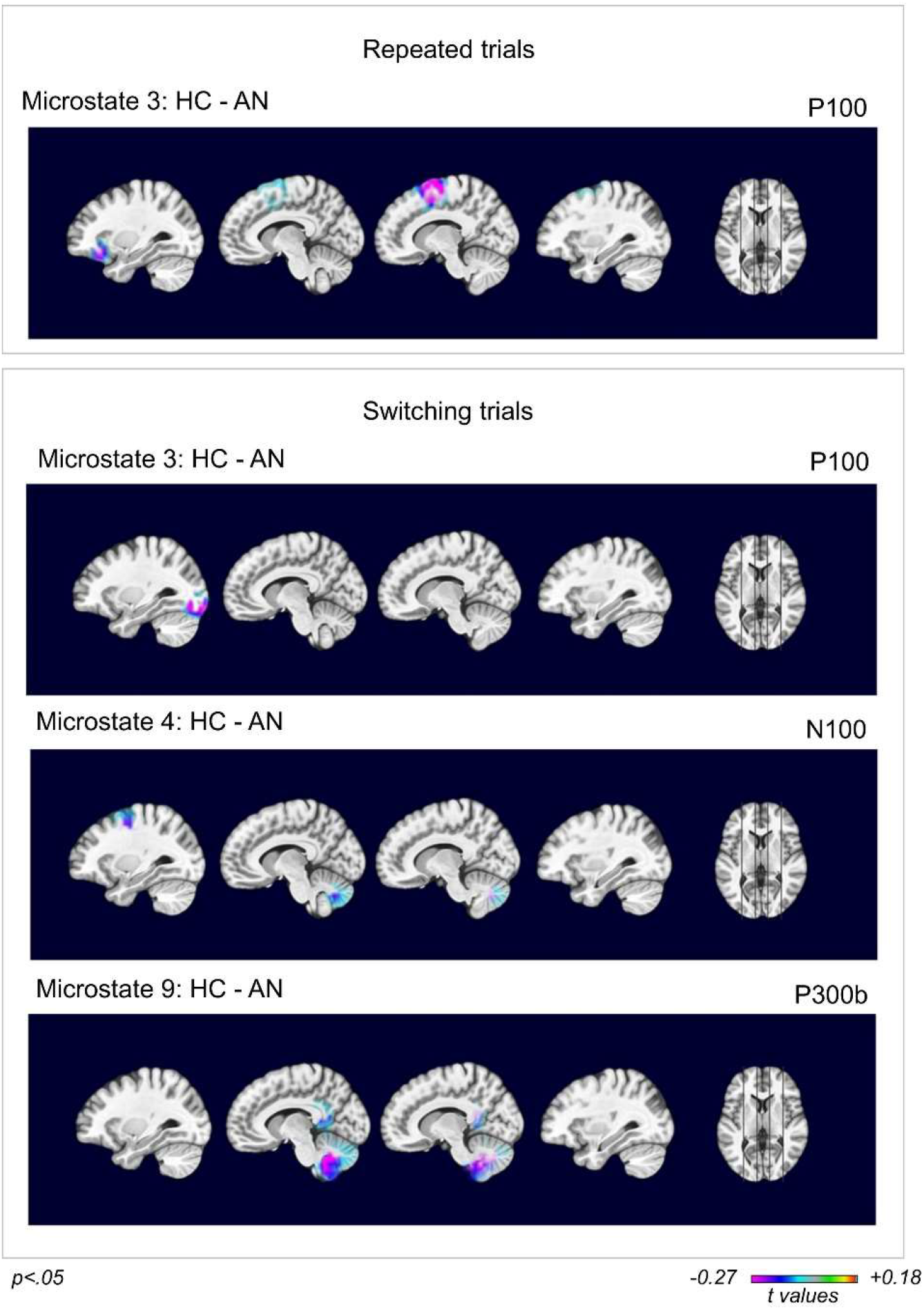
Brain imaging results. Randomization tests were performed to compare brain activity between groups. Statistically significant effects are shown for the P100 of repeated trials, for the P100, N100, and P300b of switching trials. *T* values indicate the directions of the contrast: purple/blue > activity in healthy controls (HC), green/red > activity in adolescents with anorexia nervosa (AN). The significance level was set to *p* <.05 (*p* values are corrected on regions of interest, and on average temporal windows).

**Figure 5.**
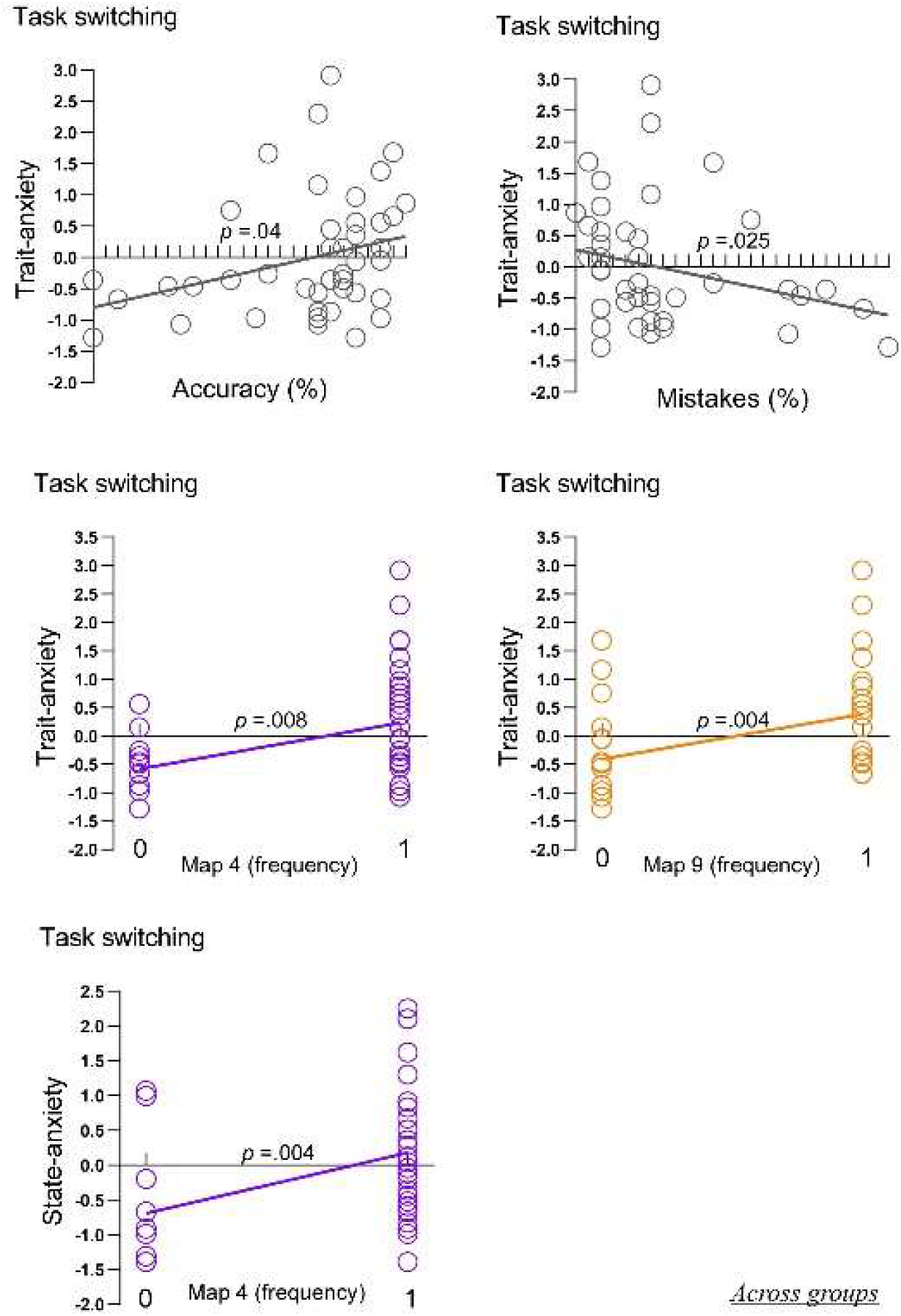
Results of correlation analysis across groups. Significant associations are shown between trait anxiety and task switching behavioral measures (p <.05), and between trait-state anxiety and microstates frequency of occurrence (p <.01).

Between-group differences in frequency of occurrence were analyzed using Pearson χ2 tests, and adjusted standardized residuals for testing cells significance. To minimize the number of multiple comparisons χ2 tests were performed on maps that were identified as potentially different between groups (i.e., for repeated trials: map 2, 3, 6, 7, 9, 10, 11; for switching trials: map 2, 3, 4, 5, 6, 7, 9, 10). For repeated trials, Map 3 differentiated the two groups [χ2(1) = 5.002, *df*= 1, *p*= .025], with higher frequency of occurrence of microstate 3 in adolescents with AN than HC (*p*=.03). No statistically significant differences were found for other maps [all χ2(1)< 1.93, *p*_*s*_> 1.6].

For switching trials, Map 3 [χ2(1) = 3.813, *df*= 1, *p*= .05] and Map 4 [χ2(1) = 4.874, *df*= 1, *p*= .027] differentiated the two groups. Adolescents with AN had significantly higher frequency of occurrence on map 3 (*p*=.03) and on map 4 (*p*=.03) compared to HC.

For all maps, group comparisons of maps’ offset, and AUC were carried out using Mann–Whitney U test. The resulting *p*-values were corrected using false discovery rate (FDR) with *p* < .05(Benjamini & Yekutieli, 2001).

For repeated trials, no significant group differences were identified on maps first offset (all *p*_*s*_ >.05). For switching trials, group differences were confirmed for offset of map 9, with the AN group showing higher values than HC [Mann–Whitney *U*= 251, *Z*=2.63, *p=*.007]. No other significant effects were found (all *p*_*s*_ >.05).

For the AUC, group differences were confirmed for map 3 of repeated trials, with adolescents with AN showing significantly higher values than HC [Mann–Whitney *U*= 339, *Z*=2.09, *p=*.036], and for map 4 of switching trials, with the AN group showing significantly higher map 4 values than HC [Mann– Whitney *U*= 349, *Z*=2.20, *p=*.027]. No other significant effects were found (*p*>.05).

### 3.4 Brain imaging results

Microstates analysis highlighted differences between groups at the P100 (map 3) for repeated trials, and at the P100 (map 3), N200 (map 4), and P300b (map 9) for switching trials. Between group analyses were performed focusing on these components, and time windows for contrast analyses were defined by k-means clustering on the grand averages.

For the P100, group differences were confirmed in both conditions (time window: 60-128 ms; microstate 3 (AN)). For repeated trials, the randomization test showed differences between AN and HC in the left fronto-orbital cortex and in the right supplementary motor area (all *p*_s_ <.05). Reduced activations were found for adolescents with AN compared to HC in all these brain regions [left fronto-orbital cortex (*t*= -2.074, *p*=.043); right supplementary motor area (*t*=-2.302, *p*=.026)]. For switching trials, the randomization test showed differences between AN and HC in the left inferior occipital cortex (*p*<.05), with reduced activations in adolescents with AN compared with HC [left inferior occipital cortex (*t*=- 2.443, *p*=0.018)].

For the N100 (time window: 136-176 ms, microstate 4 (AN)), the randomization test showed differences between AN and HC in the left precentral gyrus and in the vermis (all *p*_s_<.05). Diminished activations were found for adolescents with AN compared with HC in all these brain regions [left precentral gyrus (*t*= -2.31, *p*=.02); vermis (*t*= -2.57, *p*=.008)].

For the P300b (time window: 368-452, microstate 9 (AN)), the randomization test showed differences between AN and HC in the left and right posterior cingulate, in the left and right cerebellum, and in the vermis (p<.05). These analyses showed diminished activations in AN compared to HC in all these brain regions [left posterior cingulate (*t*= -2.07, *p*=.044); right posterior cingulate (*t*= -2.37, *p*=.02); left cerebellum (*t*= -2.23, *p*=.031); right cerebellum (*t*= -2.74, *p*=.008); vermis (*t*= -2.18, *p*=.035)].

### 3.5 Correlation analysis

As there were no significant group differences on behavioral measures of task switching, correlations analyses with trait/state anxiety were conducted across the sample as a whole. This analysis highlighted a significant positive correlation between accuracy of switching trials and trait-anxiety across groups (*r*_*pb*_=.38, *p*=.04, [.10, .50]) and a negative correlation between mistakes in switching trials and trait-anxiety across groups (*r*_*pb*_=-.35, *p*=.025, [-.59, -.05]). No other statistically significant correlations were found (*p*_*s*_>.05).

Within the AN sample, no significant correlations were found between frequency of occurrence of microstates 3 (repeated and switching trials), 4 (switching trials), and 9 (switching trials), BMI, and trait/state anxiety (*p*>.05). For the EDE-q, a significant negative correlation was found between ‘Eating concern’ and map 3 of ‘repeated trials’(*r*_*pb*_=-.43, *p*=.04, [-.78, -.02]). No other effects were statistically significant.

Further point-biserial correlations were applied to determine the associations between scores of trait-state anxiety and the presence of microstates 3 (repeated and switching trials), 4 (switching trials), and 9 (switching trials) across groups. For switching trials only, positive correlations were identified between trait-anxiety and microstate 4 (*r*_*pb*_=.38, *p*=.008, [.19, .57]) and 9 (*r*_*pb*_=4.2, *p*=.004,[.17, .63]), and state-anxiety and microstates 3 (*r*_*pb*_=.3, *p*=.04, [0.13, .55]), 4 (*r*_*pb*_=.38, *p*=.007, [.12, .61]), and 9 (*r*_*pb*_=.36, *p*=.016, [.09, .57]). On statistically significant results, a Bonferroni correction was applied to protect against false positives. After Bonferroni correction, correlations between trait-anxiety and microstate 4 and 9, and between state anxiety and microstates 4 remained significant (*p* <.0083).

### 3.6 Post-hoc tests

In order to assess whether comorbidities could have driven microstates effects on mental switching, further post hot tests were performed by removing the 3 adolescents with AN who had a disagnosis of anxiety disorder and depression. Post hoc comparisons were performed between 19 AN patients and 23 HC.

The results of this analysis confirmed group differences on Map 4 of switching trials [χ2(1) = 5.05, df= 1, *p*= .025], with higher presence of microstate 4 in adolescents with AN than HC (p=.02). The higher frequency of map 3 in AN than controls was not confirmed by this post-hoc test [χ2(1) <2.29, *p*> 1.3]. Group differences were confirmed for offset of map 9, with the AN group showing higher values than HC [Mann–Whitney *U*= 197, *Z*=2.84, *p=*.003].

## 4. Discussion

To our knowledge, this is the first high-density EEG study to investigate CF in adolescents with AN. Findings of this study suggest that neural inefficiencies of task-switching are evident in adolescents with AN. Our results indicate that AN is associated with altered early visual orienting processing (N100), and subsequent attentional mechanisms (P300b). During task switching, reduced activations in AN were identified in occipital, pre-central, posterior cingulate, and cerebellar regions. Of note, for task switching, we found that trait and state anxiety were correlated with the occurrence of atypical cluster maps of the AN sample across groups. No behavioral deficits in task switching were observed in AN. This finding is consistent with previous studies that have investigated CF in adolescents with AN.

Regarding the behavioral performance, similarly to other studies using the DCCS (Ezekiel, Bosma, & Morton, 2013; Zelazo, 2006), we found responses on switching trials significantly less accurate, slower, and more error prone than responses on repeated trials. In line with previous evidence in adolescents with AN (Andrés-Perpiña et al., 2011; Bohon et al., 2020; Castro-Fornieles et al., 2019; Fitzpatrick, Darcy, Colborn, Gudorf, & Lock, 2012), we found no evidence of impaired encoding of task-relevant information in the AN group. Therefore, our findings seem to suggest that task-switching behavioral impairments may reflect the chronicity of the disease.

Surprisingly, an enhanced quality of performance (i.e., greater overall accuracy) was observed in the AN group. This finding seems to indicate an involvement of additional strategies in adolescents with AN (e.g. enhanced effort, increased use of resources). Several explanations are plausible. One possibility is that perfectionism in adolescents with AN leads to stronger concern over mistakes (Bulik et al., 2003; Dahlenburg, Gleaves, & Hutchinson, 2019); on the other hand, it is also possible that anxiety induced an enhanced effort on task execution(Eysenck, Derakshan, Santos, & Calvo, 2007). This might be corroborated by the correlations between switching trials (positive with accuracy, negative with mistakes) and trait-anxiety across groups.

A reasonable conclusion is that if perfectionism correlates with an anxious temperament (higher trait anxiety levels), these dimensions may potentially interfere with performance quality of CF. It remains unclear whether an alternative behavioral strategy, which is more resources demanding/effortful, may also represent a risk factor for developing of executive function deficits.

ERP were investigated using microstates analysis and source imaging. Contrary to our expectations, P100 abnormalities in AN were identified in both experimental conditions. P100 responses are thought to reflect automatic recruitment of visual spatial attention(Di Russo & Spinelli, 1999), but also cognitive modulation of sensory processing (Kaiser et al., 2020). For trials that were repeated, source imaging analysis indicated differences between AN and HC in the left orbitofrontal cortex, and in the right supplementary motor area. The orbitofrontal cortex has a key role in decision making and in sensory input integration (Schuck, Wilson, & Niv, 2018), while the supplementary motor area is involved in action control(Nachev, Wydell, O’neill, Husain, & Kennard, 2007). The observed decrease in activation could indicate that adolescents with AN were quicker to automatically orient attention to task perseveration than HC. In this regard, within the AN group, a significant correlation was observed between ‘Eating concern’ and the P100 map of repeated trials. This indicates that dysfunctional brain patterns of task perseverance may be linked to active illness symptomatology.

For switching trials, differences between AN and HC on the P100 were identified in the left inferior occipital cortex. This visual region has been previously linked with task switching(Buchsbaum, Greer, Chang, & Berman, 2005; Nyhus & Barceló, 2009), and previous literature has shown reduced activations to task switching in the inferior occipital cortex in adolescents with AN (Castro-Fornieles et al., 2019). Our findings appear consistent with these studies, and suggest a reduced sensory-perceptual sensitivity to change detection. However, effects on the P100 of switching-trials were not present once adolescents with an emotional disorder diagnosis were excluded from the analyses, highlighting that sensorial-perceptual abnormalities may be driven by full blown emotional disorder diagnosis.

During task switching, group differences were also detected for the N100. The N100 is an early sensorial orienting component(Hillyard & Anllo-Vento, 1998), that is elicited by matching processes(Sur & Sinha, 2009). Source imaging analyses indicated lower activations in the left precentral gyrus and the vermis in adolescents with AN when compared to HC. These regions are responsible for motor control(Banker & Tadi, 2019) and integration of sensory signals (Schmahmann, 2019). In adults with AN, previous studies have indicated alterations in the left precentral gyrus (Barona et al., 2019) and in cerebellar networks (Amianto et al., 2013; Castro-Fornieles et al., 2019). The findings of this study seem to highlight a decreased frontal/cerebellar functionality most probably implicated in sensory integration into a motor program.

P300 latencies are thought to be related to the speed of mental processes(Sur & Sinha, 2009). In this study, shorter latency of microstate 9 (P300b) was observed in adolescents with AN when compared to HC. For the P300b, source imaging analysis indicated reduced activations in adolescents with AN in the bilateral posterior cingulate, cerebellum, and in the vermis. The posterior cingulate cortex is deactivated during cognitive tasks involving externally directed attention (Singh and Fawcett 2008), while the cerebellum, among other cognitive regulatory functions, is also involved in set-shifting, and perseveration (Schmahmann, 2019). A prematurely modulated attentional/executive network, likely to involve additional neural resources for task switching, is suggestive of inefficient neural strategies in AN.

Dysfunctional N100 and P300 b microstates were predicted by anxiety scores across groups. For switching trials, correlation analyses indicated a positive association between trait-anxiety and presence of microstate 4 and 9, and between state-anxiety and microstate 9. These findings suggest that high levels of trait-state anxiety may account for dysfunctional neural strategies typical of AN. Consequently, although studies in AN adults seem to rule out a link between anxiety symptoms and CF dysfunctions, this study suggests that trait anxiety may explain neural profiles that are characteristic of AN. Correlation analyses within the AN group demonstrated that neural inefficiencies of switching trials were independent of active symptomatology and BMI. This indicates that dysfunctional neural patterns of task switching were independent of the active symptomatology of the illness.

This study has some limitations. The main limitation is the relatively small sample size, which could have limited the statistical power of our analysis. The limited samples size did not allow to investigate subtypes of AN. Furthermore, one participant taking psychotropic medication was included: however, we did assess the possible influence of comorbidities. Further studies should include larger sample size in order to replicate our findings.

This study sheds novel insights into potential mechanisms mediating behavioral characteristics and neural vulnerabilities for mental inflexibility in adolescents with AN. Our work suggests inefficient behavioral and neural strategies in adolescent with AN. Behavioral performance on a CF task seems to indicate an enhanced cognitive effort on task execution in the AN group. Neural abnormalities of the N100 and of the P300 b might be characteristic of adolescent AN during task switching: these are localized in large-scale brain networks including the left precentral gyrus, posterior cingulate, and cerebellum. Even in the absence of behavioral impairments, network inefficiencies may precede behavioral deficits associated with CF in subsequent stages of the illness. Our findings also indicate that abnormal neural patterns characteristic of AN might be related to trait and state anxiety. Insights generated by this study increase our neuro-biological understanding of adolescent AN, as well as the role of anxiety in CF neural and behavioral traits in AN.

## Acknowledgements

This work was supported by the Foundation Gertrude von Meissner and the *Fondation Ernst et Lucie Schmidheiny* (to CB, and NM).

